# Visual field map size constrains working memory precision

**DOI:** 10.64898/2026.06.01.729356

**Authors:** Nathan Tardiff, Xingyu Ding, Xiao-Jing Wang, Clayton E. Curtis

## Abstract

Individual differences in working memory are associated with cognitive functioning and real-world outcomes, including general intelligence, academic achievement, and psychiatric and neurological disorders. Despite its importance, the neurobiological mechanisms that drive individual differences in working memory are unknown. Here, we asked whether the precision of spatial working memory covaries with the cortical surface area of retinotopically organized maps that are thought to store the content of working memory. In particular, we hypothesized that individuals with larger visual field maps would have more precise working memory. We assessed human subjects’ (male and female) working memory precision using a memory-guided saccade task, and using fMRI we separately mapped retinotopically organized regions of visual, parietal, and frontal cortex. In support of our hypothesis, we found that people with larger maps in primary visual cortex and parietal cortex had better memory. To test hypotheses about the potential mechanisms driving our results, we simulated working memory in neural network models with varying numbers of synthetic neurons. Echoing our empirical results, larger networks exhibited less memory error. This reduction in error could be fully attributed to a decrease in the proportion of memory-perturbing noise along the attractor in larger networks, establishing a pure size effect on working memory precision.

## INTRODUCTION

Whether navigating bustling city streets or overcrowded computer displays, we often need to hold and manipulate information in mind that is no longer available via perception. This cognitive capacity, termed working memory, is critical to cognitive functioning (Tardiff & Curtis, 2025). Without it, our cognition is bound to the present, and “out of sight is out of mind” (Goldman-Rakic, 1995). Working memory varies considerably across individuals, and this variation is associated with many other cognitive functions and life outcomes (Barrett et al., 2004; Johnson et al., 2013; Luck & Vogel, 2013; Unsworth & Engle, 2007). Individual variation in working memory predicts general intelligence (Conway et al., 2003; Engle et al., 1999; Fukuda et al., 2010; Shipstead et al., 2012; Süß et al., 2002) and is correlated with broader executive functions (Miyake et al., 2001). Working memory is important for reading and linguistic comprehension (Daneman & Carpenter, 1980; Hasher & Zacks, 1988), and academic achievement (Alloway, 2006; Alloway et al., 2009; Ashkenazi et al., 2013). Working memory declines with aging (Hasher & Zacks, 1988; Salthouse et al., 1991), and working memory deficits are a characteristic feature of many psychiatric conditions, including schizophrenia (Barch, 2005; Gold et al., 2006; Johnson et al., 2013; J. Lee & Park, 2005; Park & Holzman, 1992), Parkinson’s (E.-Y. Lee et al., 2010), and ADHD (Alderson et al., 2013; Rapport et al., 2008).

Despite its significance, the mechanisms that determine the quality of working memory across individuals remain unknown. In the domain of visual working memory, while functional neural correlates of individual variation in storage capacity and attentional control have been identified (Fukuda et al., 2015; Mitchell & Cusack, 2008; Todd & Marois, 2005; Unsworth et al., 2015; Vogel & Machizawa, 2004; Xu & Chun, 2006), we lack an understanding of the neurobiological mechanisms that underlie these differences. This in turn limits our ability to target these mechanisms in clinical interventions.

A potential constraint underlying individual differences is the amount of cortex available to support a particular function. In visual perception, the amount of early visual cortex —particularly V1—devoted to representing different parts of the visual field varies considerably, and these differences mirror differences in perceptual performance across the visual field (Benson et al., 2021; Duncan & Boynton, 2003; Harvey & Dumoulin, 2011; Himmelberg et al., 2023; Hubel & Wiesel, 1974; Silva et al., 2018). Furthermore, individual variability in the size of early visual cortex predicts individual differences in positional acuity (Duncan & Boynton, 2003; Silva et al., 2021; Song et al., 2015), contrast sensitivity (Himmelberg et al., 2022; Jigo et al., 2023), orientation discrimination (Song et al., 2013), motion perception (Murray et al., 2024), and the strength of context effects (Schwarzkopf et al., 2011; Song et al., 2013). These results suggest perception is impacted by aspects of neural architecture that covary with size, such as the overall number of neurons and/or the connectivity and response properties of neurons, generally leading to better performance with larger visual areas (Song et al., 2015). Visual working memory is thought to be subserved in part by the same cortical areas that underlie perception (Curtis & D’Esposito, 2003; D’Esposito & Postle, 2015; Serences, 2016). Supporting this idea, the contents of working memory can be decoded and reconstructed from several retinotopically organized visual field maps spanning visual, parietal, and frontal cortex (Christophel et al., 2017; Ester et al., 2015; Hallenbeck et al., 2021; Harrison & Tong, 2009; Kwak & Curtis, 2022; Li et al., 2021, 2025; Rademaker et al., 2019; Serences et al., 2009; Tardiff & Curtis, 2025).

Given the cortical overlap between perception and working memory, in this study, we test the hypothesis that the precision of visual working memory will be greater in individuals with larger visual field maps. To gain mechanistic insight into why larger maps might lead to greater precision, we simulated working memory in artificial neural network models of varying size. While most prior work on individual differences in visual working memory has focused on measures of working memory capacity (i.e., the number of items that can be stored, e.g., (Luck & Vogel, 2013), we focus on the precision with which memoranda can be recalled for two reasons. First, precision likely provides a purer and more accurate measure of the fidelity with which the neural substrates of storage can encode and maintain information, whereas measuring capacity limits commonly involves additional demands on attentional control processes (Fukuda et al., 2015; Fukuda & Vogel, 2009, 2011; Shipstead et al., 2014) and procedurally often lacks a continuous measure of working memory recall. Notably, memory factors—not attentional factors—predict general intelligence (Shipstead et al., 2014). Second, such continuous measures of precision can be more tightly linked to measures of precision derived from the activity of neural network models.

To test our hypothesis, we used an fMRI-based retinotopic mapping procedure to precisely define visual field maps in individual subjects, and we then related the surface area of these maps to the precision of subjects’ visual working memory, as measured by a memory-guided saccade task. To preview, we found that the sizes of select maps in visual and parietal cortex are correlated with working memory precision. Moreover, using our network model we identified a potential mechanism underlying this size effect, whereby increasing size decreases the degree to which noise in the network can perturb its memory representation.

## RESULTS

### Strong interindividual variability in retinotopic map sizes

We focused our analyses on five retinotopically organized regions of interest (ROIs) along the dorsal stream that have been implicated in visual working memory: two ROIs in visual cortex (V1, V3AB), two along the intraparietal sulcus (IPS0/1, IPS2/3), and one in frontal cortex in the superior portion of the precentral sulcus (sPCS). In each subject (N = 24), we fit a population receptive field model to the BOLD data from a retinotopic-mapping task to generate polar-angle and eccentricity maps across visually responsive cortex (Kay et al., 2013; Mackey et al., 2017); Figure 1A). ROIs were then delineated using established functional and anatomical criteria, where boundaries between adjacent areas lie along polar-angle reversals (Mackey et al., 2017; Silver & Kastner, 2009). We then established the size of each ROI by measuring its surface area.

**Figure 1:**
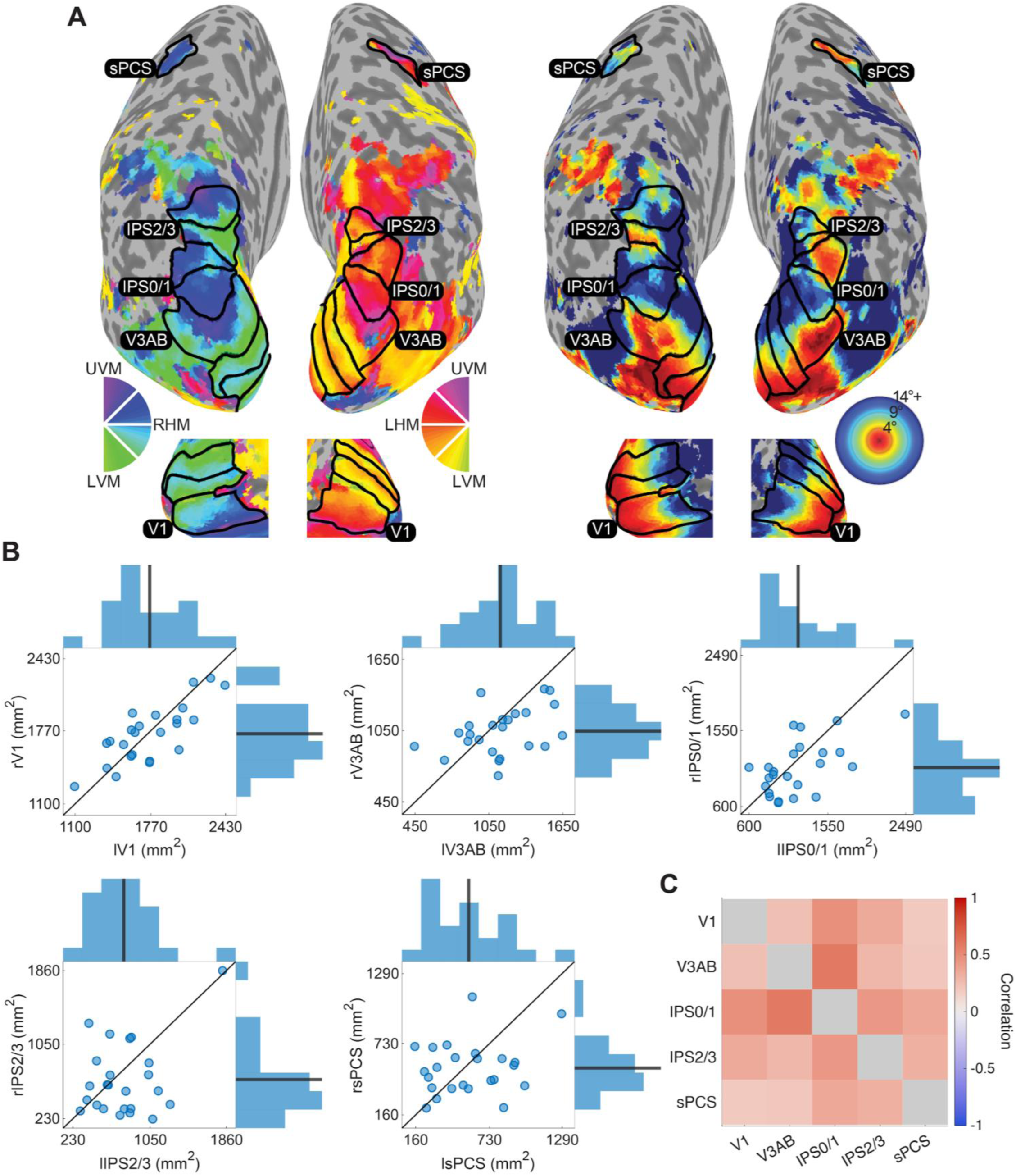
Map sizes vary substantially across individuals, hemispheres, and ROIs. **A** ROI definitions are outlined on the cortical surface for an example subject. ROIs were identified at the single-subject level using population receptive field mapping (Mackey et al., 2017), which yields polar angle (left) and eccentricity (right) maps across visually responsive cortex. The maps shown are thresholded at ≥ 10% variance explained at each vertex. Only ROIs included in the analyses are labeled. Note that IPS0/1 and IPS2/3 are composites of pairs of IPS ROIs. **B** The relationship between left hemisphere (abscissa) and right hemisphere (ordinate) surface area is plotted for each ROI. Points are surface-area values for individual subjects, and lines are the unity line. Marginal histograms for the ROI in each hemisphere are along the corresponding axis, where the thick line is the mean surface area across subjects. See Figure 1 - Supplement 1 for unaggregated IPS results. **C** Intercorrelations among ROI sizes were weak to moderate. Correlations were computed with data collapsed across hemispheres (i.e., each ROI has two values per subject), using Spearman’s *ρ*.

**Figure 1 - Supplement 1:**
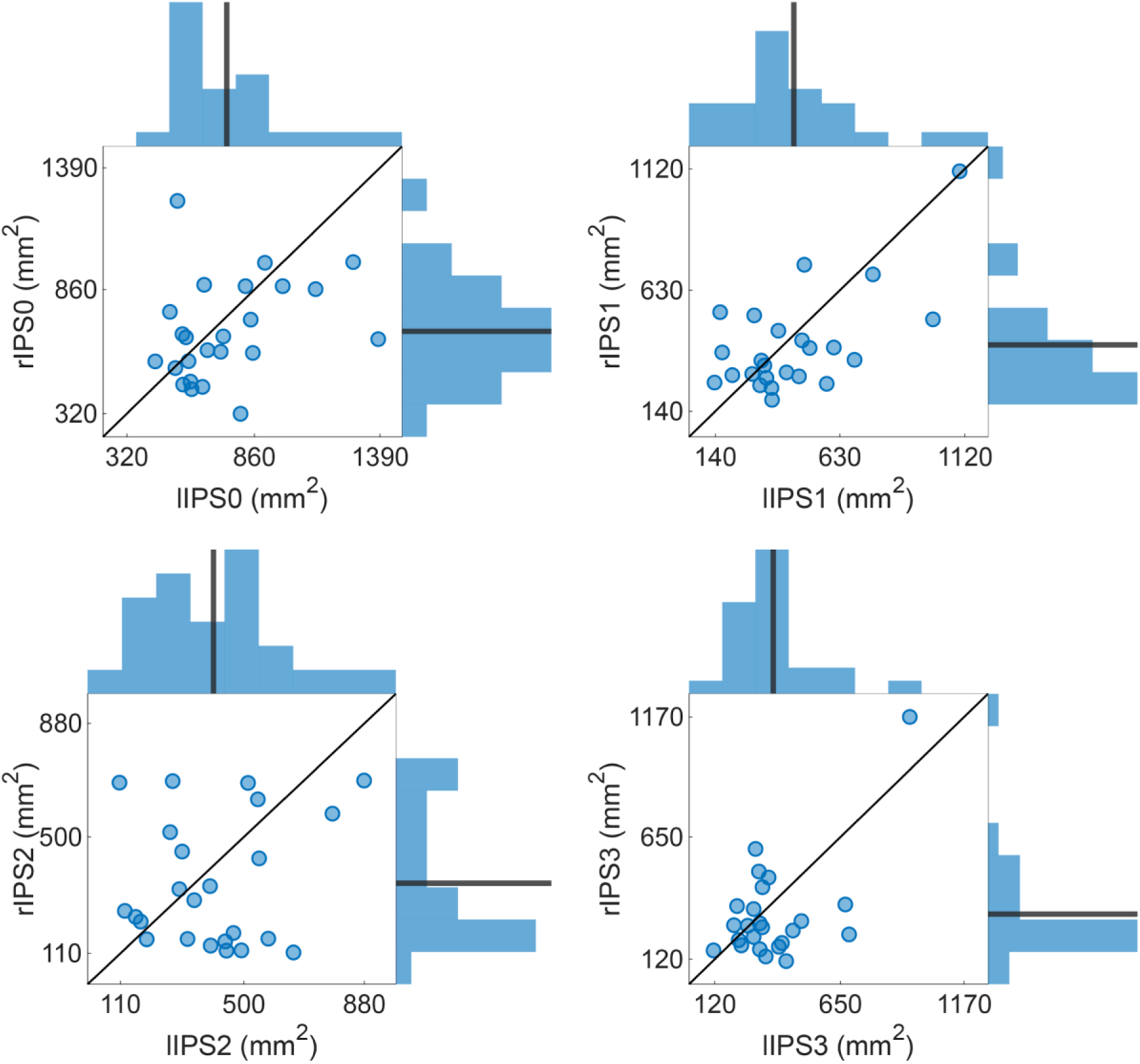
IPS ROI variability. The relationship between left hemisphere (abscissa) and right hemisphere (ordinate) surface area is plotted for unaggregated IPS ROIs. Points are surface-area values for individual subjects, and lines are the unity line. Marginal histograms for the ROI in each hemisphere are along the corresponding axis, where the thick line is the mean surface area.

We characterized the variability within and between ROIs across subjects and hemispheres before asking whether individual differences in map size are associated with working memory. The size of V1 varied approximately 2-fold across individuals (coefficient of variation (CV) = 0.167), and was strongly correlated between hemispheres (Figure 1B; *ρ* = 0.73, *p_FDR_* = 0.001), replicating previous reports (Andrews et al., 1997; Benson et al., 2022; Dougherty et al., 2003; Harvey & Dumoulin, 2011). Size variability increased up the visual hierarchy, peaking at IPS2/3 (CV = 0.522) and was only slightly less in sPCS (CV = 0.462), both corresponding to over 8-fold variation across subjects. Unlike V1, correlations between hemispheres were weak to moderate for most other ROIs, only reaching significance in IPS0/1 (*ρ* = 0.60, *p_FDR_* = 0.01). Size variability was not driven by gross anatomical differences such as brain size or biological sex. Variability in every ROI was larger than variability in total cortical surface area (CV = 0.07). While males had larger total cortical surface area on average than females (M_diff_ = 17877 mm^2^, *t*(21.975) = 4.66, *p* < 0.001)—in line with established anatomical differences (Eliot et al., 2021; Williams et al., 2021)—the sizes of individual ROIs did not vary as a function of sex (all *p*s*_FDR_* > 0.05). Further indicating that each ROI could be a source of independent variance in behavior, ROIs were only weakly to moderately correlated with each other (Figure 1C).

### Working memory precision varies across individuals

To assess visual working memory, subjects completed a memory-guided saccade task (Figure 2A). Two visual targets were presented briefly —one per hemifield—and after a delay, subjects were cued to report the location of one of the targets by moving their eyes to the remembered position.

**Figure 2:**
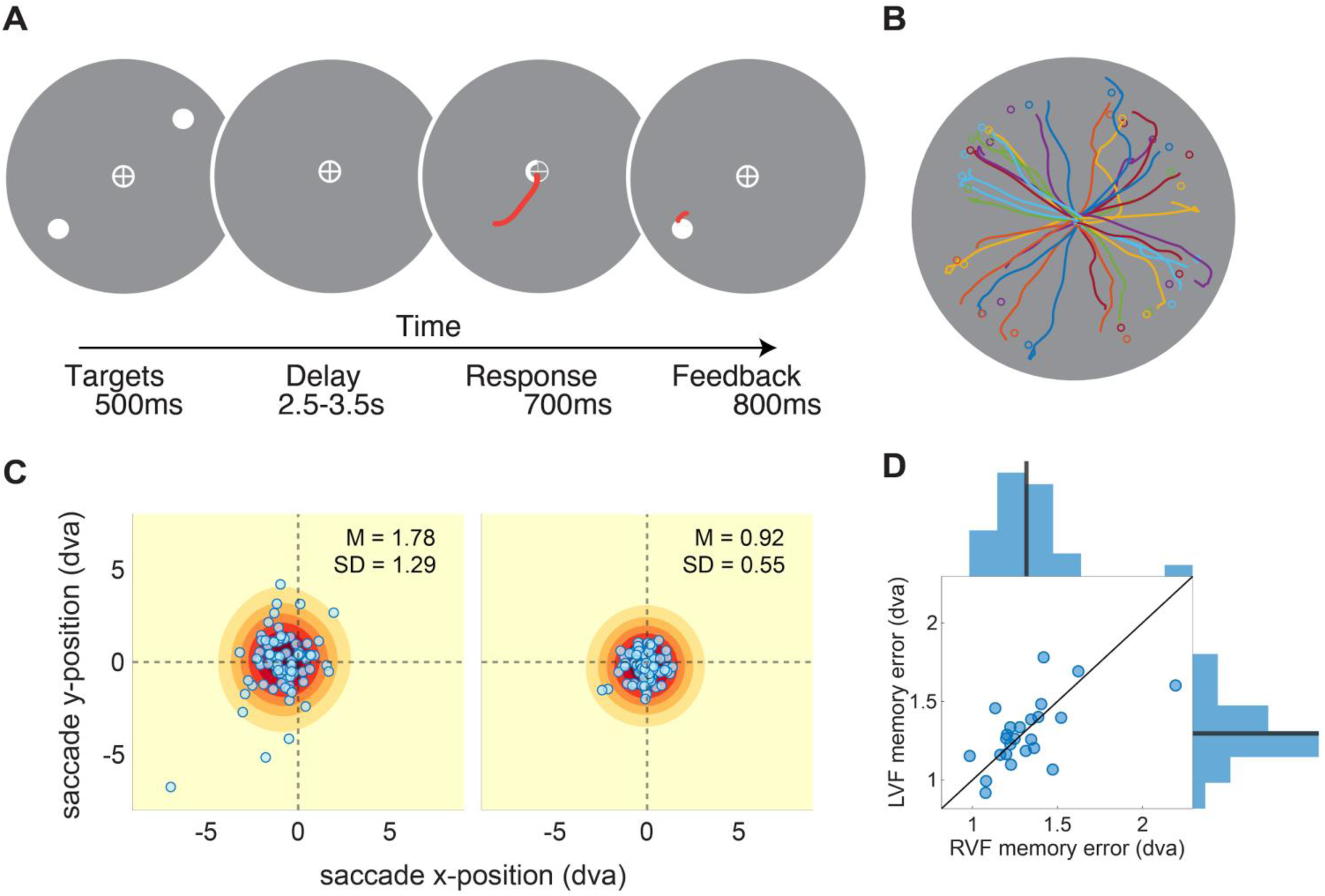
Working memory precision varies across subjects. **A** Memory-guided saccade task. Two items were presented, one in each hemifield. After a delay, a response cue instructed which item should be reported with a memory-guided saccade. Feedback was provided, followed by an intertrial interval (2,000-3,000 ms; see Methods). **B** Saccade trajectories (colored lines) and true item locations (circles) from an example task run for one subject. Memory errors were calculated as the Euclidean distance between the final saccade endpoints and the true locations. **C** Memory errors from the left visual field for a subject with higher error (left) and a subject with lower error (right). All trials were rotated such that the true item is at the origin, rightward of fixation. Dots are saccade endpoints of individual trials, and contours depict the empirical distribution over these endpoints (kernel density estimate). **D** The relationship between right visual field (left hemisphere; abscissa) and left visual field (right hemisphere; ordinate) memory error. Points are the mean memory error of individual subjects, and the line is the unity line. Marginal histograms for the error in each hemifield are along the corresponding axis, where the thick line is the mean error across subjects.

Working memory reports were relatively precise on average, with variability in the magnitude of memory error across subjects (Figure 2C,D; M (SD) right visual field: 1.318 (0.239) degrees visual angle (dva); left visual field: 1.296 (0.208) dva). Memory error was moderately correlated across hemifields (*ρ* = 0.59, *p* = 0.003), but with no systematic difference in performance between hemifields (*t*(23) = 0.527, *p* > 0.05). Given this result and the generally low to moderate correlations between map sizes across hemispheres, we next assessed the relationship between map size and working memory across hemispheres, rather than aggregating hemispheres within subject (see (Harvey & Dumoulin, 2011; Silva et al., 2021) for a similar approach).

### Select map sizes impact working memory

To ask whether the size of any of the visual field maps was associated with working memory, we computed zero-order and semipartial Spearman’s correlations between ROI surface area and memory error in the contralateral hemifield. Only the surface area of IPS2/3 (*ρ* = −0.41, *p_FDR_* = 0.014) and V1 (*ρ* = −0.41, *p_FDR_*= 0.046) significantly correlated with memory error, while relationships between map size and error in the other ROIs were weak and not significant (Figure 3; see Figure 3 - Supplement 1 for unaggregated IPS results). These correlations were largely unaffected after partialling out the effects of hemisphere, retinotopic mapping task, and total cortical surface area, though the V1 correlation was no longer significant (IPS2/3: *ρ* = −0.39, *p_FDR_*= 0.02; V1 *ρ* = −0.36, *p_FDR_* = 0.09). Consonant with the preference for the contralateral hemifield across ROIs (Figure 1A; (Mackey et al., 2017), correlations between ROI sizes and memory error in the ipsilateral hemifield did not reach significance (all *p*s_FDR_ > 0.05). These results suggest that the relationship between size and error in V1 and IPS2/3 was driven by retinotopically organized properties of the memory representation contained in the contralateral hemisphere and not by other properties of these regions.

**Figure 3:**
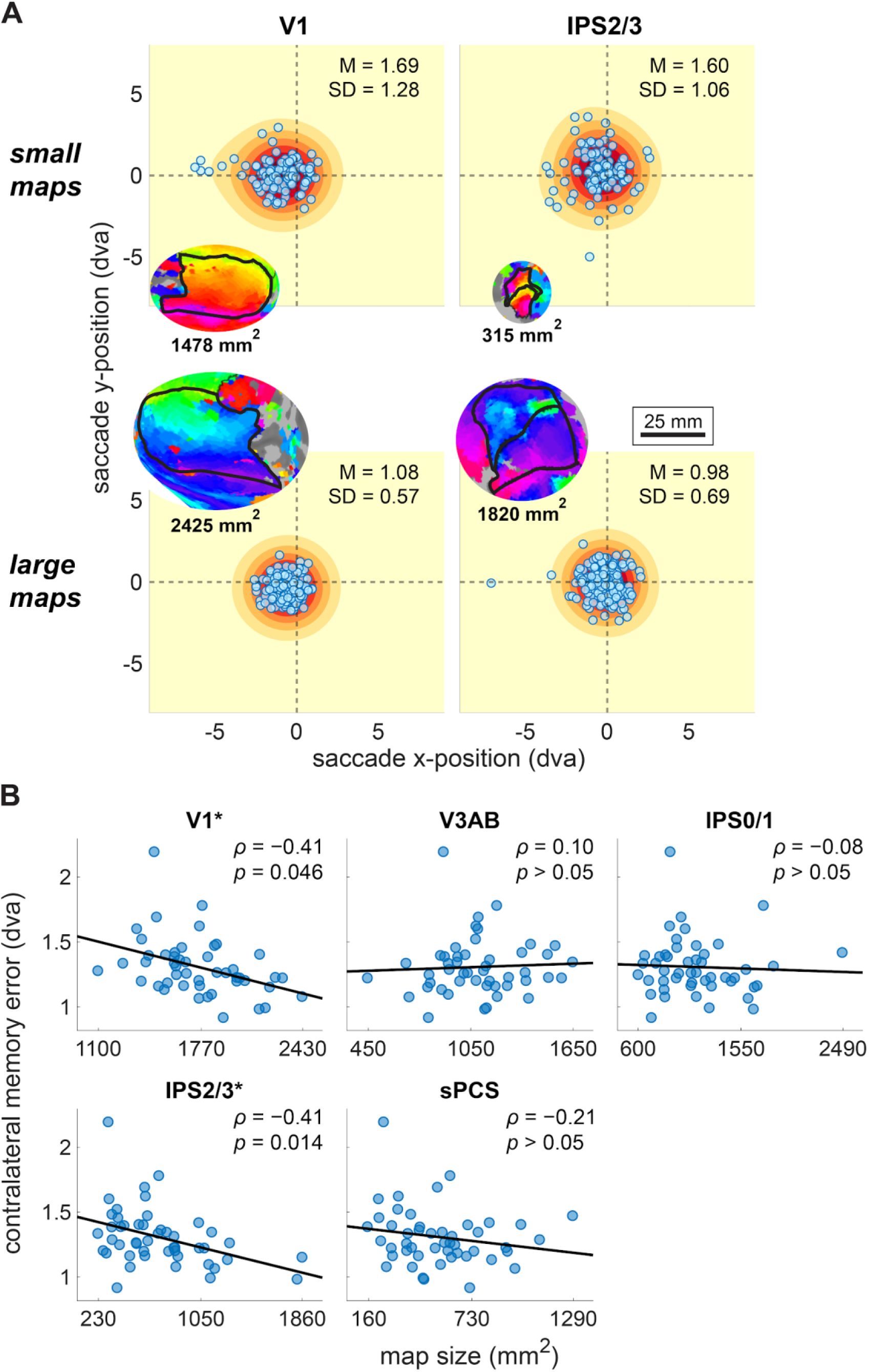
Surface area of visual field maps correlates with working memory in select areas. **A** Distributions of memory errors across all trials from the contralateral hemifield for example subjects with smaller maps (top row) and larger maps (bottom row) for V1 (left) and IPS2/3 (right). Points and contours were computed as in Figure 2C. Inset images depict the corresponding map (to scale) for each subject. **B** The relationship between map size and average contralateral memory error for all ROIs. Points are individual hemispheres from individual subjects. Linear fit lines are for illustration only; statistics were computed using Spearman’s *ρ*, and *p*-values are FDR corrected. * *p_FDR_* < 0.05

**Figure 3 - Supplement 1:**
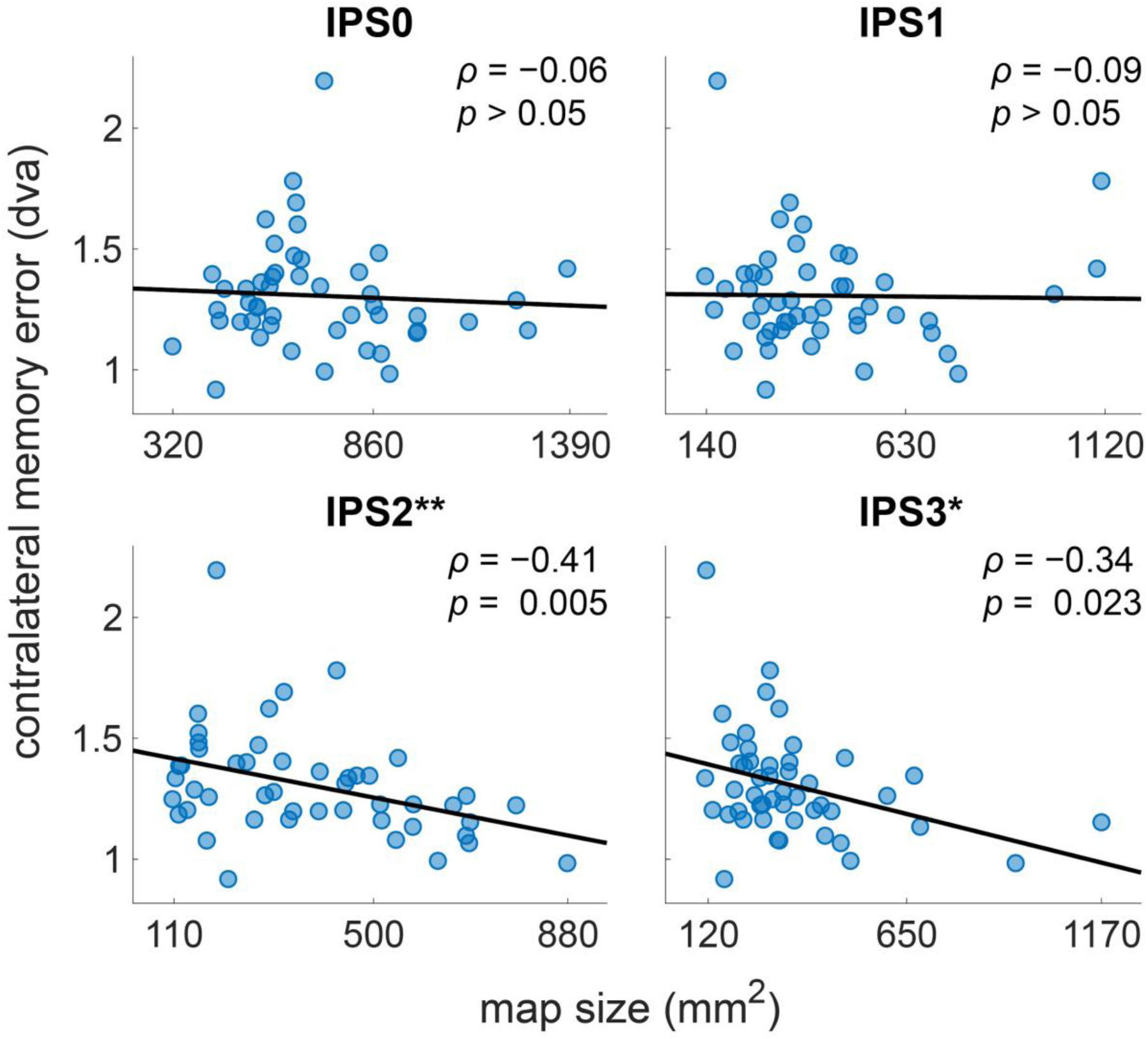
Relationships between IPS map sizes and working memory. The relationship between map size and average contralateral memory error for unaggregated IPS ROIs. IPS2 (*ρ* = −0.41, *p* = 0.005) and IPS3 (*ρ* = −0.34, *p* = 0.023) were significantly correlated with memory error individually. Points are individual hemispheres from individual subjects. Linear fit lines are for illustration only; statistics were computed using Spearman’s *ρ*, and *p-*values are uncorrected. \*\**p* < 0.01; \**p* < 0.05

In sum, though the contents of spatial working memory can be decoded from all five areas (Li et al., 2021, 2025), the correlation between map size and memory precision was specific to IPS2/3 and to a lesser extent V1, and these relationships were not readily explained away by the overall size of the cortex. This pattern of results suggests that these two areas may be particularly important for working memory storage. Somewhat surprisingly, the size of our frontal ROI, sPCS, did not correlate with performance (*ρ* = −0.21, *p_FDR_* = 0.291), despite causal evidence for its role in working memory (Hallenbeck et al., 2025; Mackey, Devinsky, Doyle, Meager, et al., 2016; Mackey & Curtis, 2017). It may be that the retinotopic structure in frontal cortex is less organized, making our estimates of the size of sPCS more challenging than those of maps in visual and parietal cortex. We return to this possible issue in the discussion. Next, we address why the size of a cortical area might impact memory precision.

### Size affects working memory error in a neural network model

To establish a mechanistic basis for the role of cortical size in working memory, we simulated a working memory task in a neural network model (Engel et al., 2015; Engel & Wang, 2011). The model features tuned recurrent excitation balanced by broad inhibition, allowing it to maintain a bump of activity encoding a stimulus angle across a delay. Each unit in the network is tuned to a particular angle, uniformly tiling the circular space and forming a ring attractor (Figure 4A), such that the memorandum can be read out by a linear population vector decoder, which weights the tuning of each unit by its firing rate (Figure 4B; (Compte et al., 2000; Georgopoulos et al., 1986; Seung & Sompolinsky, 1993). When noise is introduced into ring attractor networks, the bump of activity drifts stochastically over time, leading to memory errors of the sort found in the working memory of humans and nonhuman primates (Compte et al., 2000; Schapiro et al., 2022; Schneegans & Bays, 2018; Wimmer et al., 2014). To explore the effects of network size on decoded error, we simulated the network at four different sizes between 400 and 2000 units. Crucially, while many factors could in principle covary with size in a biological cortical network, we scaled the network connectivity such that the activity bump should be invariant to size (e.g., its spread in stimulus space and max firing rate), thereby reducing the influence of factors other than the number of units.

**Figure 4:**
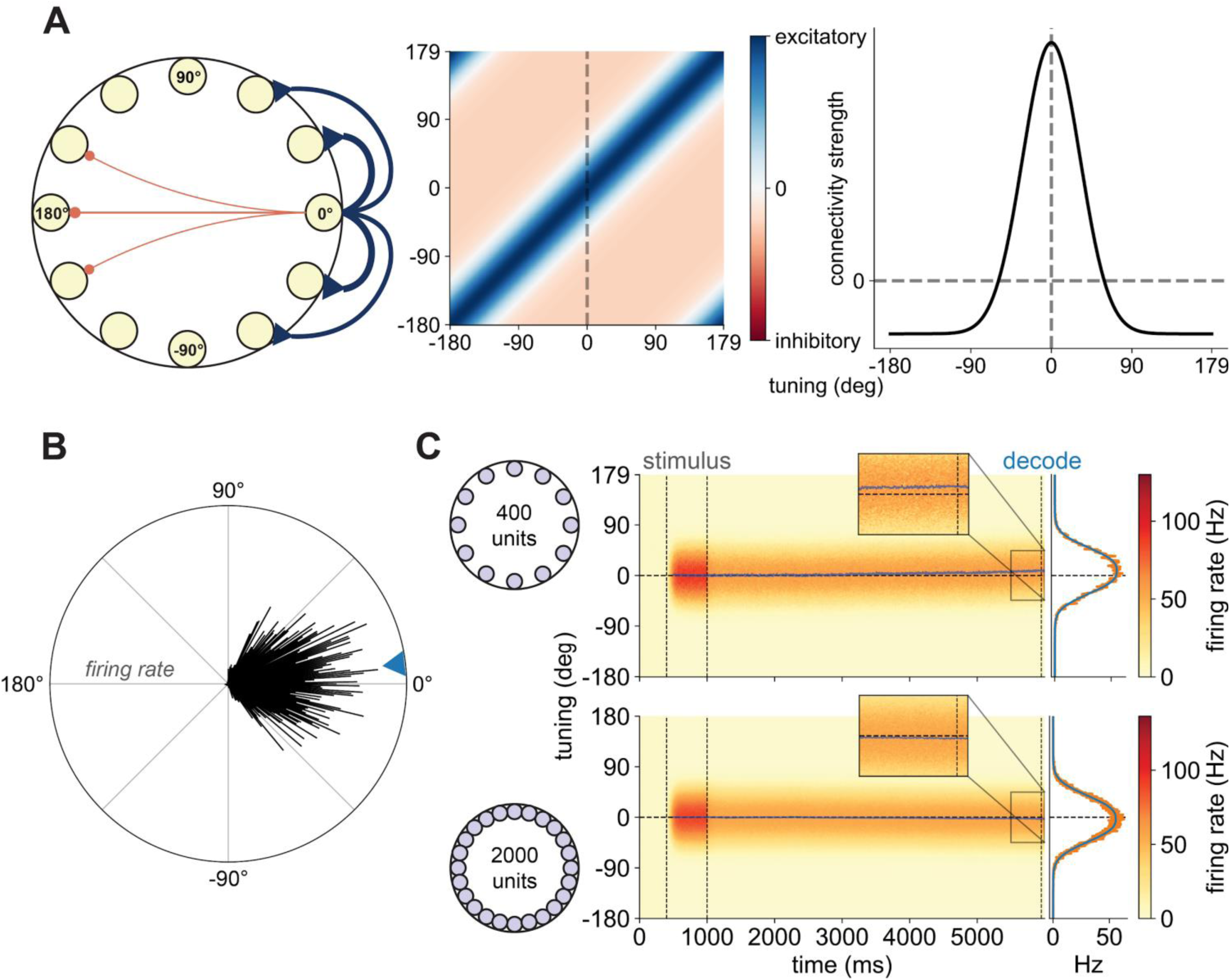
A neural network model of working memory. **A** The ring attractor model can simulate working memory via recurrent excitation coupled with broad inhibition (Engel et al., 2015; Engel & Wang, 2011). Connectivity was structured based on tuning similarity, such that units with more similar tuning have stronger excitatory connections and units with more dissimilar tuning have (weakly) inhibitory connections (left). The connectivity matrix for the network (center), where connectivity strength and sign (excitatory or inhibitory) are indicated by color. Note that the matrix for the 400-unit network is shown, but connectivity for the larger networks will appear the same due to scale-invariance. The connectivity profile for a single unit tuned to 0° (right, corresponding to the dashed line in the connectivity matrix). Note that connectivity is structured according to a Gaussian connectivity profile. **B** The population vector method for decoding the memorandum.. Black lines are vectors depicting the firing rate at the last time point of the delay of every unit from a simulation of the 400-unit network (see Figure 4C). Vector magnitudes indicate the firing rate of each unit and vector polar angles the preferred tuning. The decoded angle (blue triangle) is the polar angle of the mean firing rate vector (i.e. an average of the individual unit tunings, weighted by their firing rates). **C** Example simulated trials from the 400-unit network (top) and the 2000-unit network (bottom), with units on the ordinate, time on the abscissa, and color indicating firing rate. After stimulus presentation, the network maintained a noisy memory of the stimulus in a bump of persistent activity. This bump can drift over time, leading to memory error. The blue line is the decoded position at each time point of the delay. Inset: the bump during the last 500 ms of the delay. Error analyses averaged decoding over the last 50 ms of the delay (dashed line). Right panel: 1D depiction of the bump, computed from the averaged firing rate over the last 50 ms of the delay (orange) and fit with a generalized Gaussian function (blue).

Figure 4C shows typical simulations from the 400- and 2000-unit networks. As expected, in both networks, the bump of activity is maintained across the delay after stimulus offset, and the bump profile appears invariant to network size. While both networks maintained a fairly faithful representation of the stimulus presented at 0°, the smaller network demonstrated more drift in the decoded angle across the delay. In line with these examples, across 100 runs at each size, we found that average decoded error at the end of the delay monotonically decreased with network size (Figure 5A, solid line), with the difference in error distributions between 400- and 2000-unit networks qualitatively matching the difference between a subject with a small IPS2/3 and one with a large IPS2/3 (Figure 5B).

**Figure 5:**
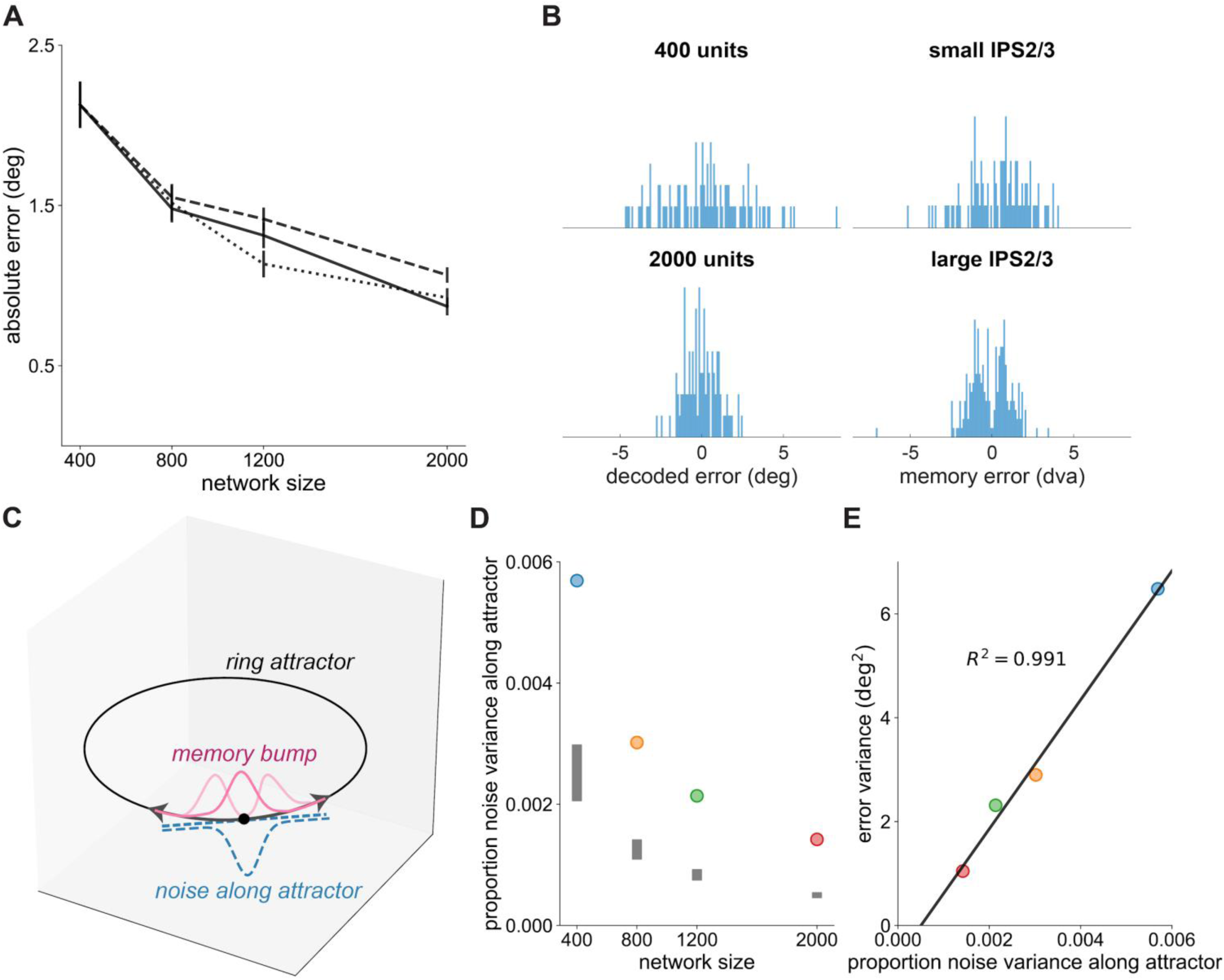
Size decreases memory error in a neural network model by reducing noise along the attractor. **A** Absolute error as a function of network size. Absolute error was defined as the unsigned difference between the stimulus at 0° and the decoded memorandum, averaged over the last 50ms of the delay. Solid line: error when decoding from all units at every size; dashed: error when decoding from the 400 units at each size matched in tuning to the 400-unit network; dotted: error from alternative simulations using networks constructed by duplicating the units in the 400-unit network *k* times to reach a given network size. Error bars are SEM. **B** Qualitative similarity between size effects in networks and humans. Histograms of the decoded error for all runs of the 400- and 2000-unit networks (left) and the memory error data of the human subject with a smaller IPS2/3 and one with a larger IPS2/3 from Figure 3A (right). Decoded error in the networks was computed as the average of the signed error in the last 50 ms of the delay. Human memory error was reduced to one dimension as the Euclidean error signed by whether the saccade was above (positive) or below (negative) the target. **C** Schematic of the interaction between the working-memory bump and network noise. Lighter pink memory bumps depict possible drifts of the bump at later time points due to the effect of noise along the attractor. See Methods for details. See (Moreno-Bote et al., 2014) for a similar illustration of the related phenomenon of information-limiting correlations in perception. **D** The proportion of total noise variance along the attractor, as a function of network size. Points are the proportion of variance. Shaded regions demarcate the 2.5–97.5 percentiles of the null distribution formed by finding the proportion of noise variance along random vectors. Note that the proportion of variance along the attractor decreases with size but is consistently above that expected for a random vector. **E** The variance of memory errors at the end of the delay as a function of the proportion of noise variance along the attractor for every network size. Error variance was calculated over the signed decoded error, averaged over the last 50 ms of the delay. Points are colored by network size as in **D.** The line is a linear fit to the data.

**Figure 5 - Supplement 1:**
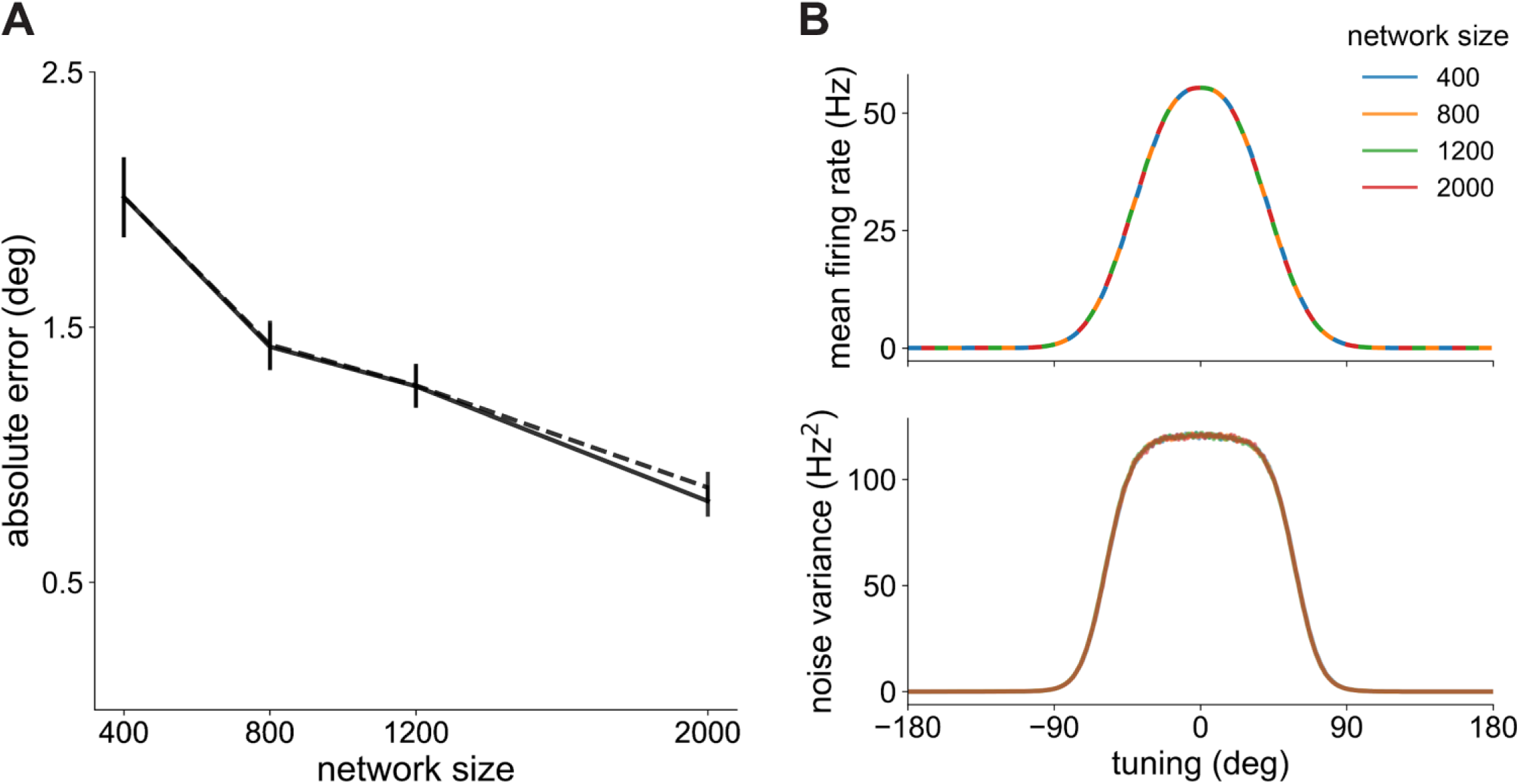
Network error (alternative measure), mean, and variance. **A** Absolute error as a function of network size. Error was defined as the absolute value of the averaged signed difference between the stimulus at 0° and the decoded memorandum over the last 50 ms of the delay. Lines are the error for the standard networks (solid) and standard networks only decoding from 400 units at each size (dashed). Error bars are SEM. **B** Mean bump profile (top) for all network sizes (colors; see legend). Lines for each size are offset to show the overlap across sizes. To generate the mean bump profile at each size, generalized Gaussian parameters fit to the mean activity in each 50 ms time window (*t* > 2000 ms) were averaged together (Supplementary Table 1), except for the *μ* parameter determining bump position, which was set to zero. Residual noise variance (bottom) at each size is also almost completely overlapping. Noise variance was computed over the residual activity after subtracting the grand mean bump profile in that time window, which was computed using the fit *μ* for that time window and the mean values for all other parameters, averaged across time points and sizes. Noise variance was computed within each run and then averaged across runs. Error bars (largely not visible) are SEM.

**Supplementary Table 1:** Fit bump parameters. Average parameters from generalized Gaussian functions fit to the mean firing rates at each network size *N* across 50 ms non-overlapping time windows. These parameters, which control the shape and height of the function fit to the activity bump, are averaged within each condition and then across conditions (grand mean). See Methods for parameter descriptions and fitting details.

| $N$ | $r_{min}$ | $g_b$ | $\alpha$ | $\beta$ |
| --- | --- | --- | --- | --- |
| 400 | 0.046 | 55.351 | 50.380 | 2.522 |
| 800 | 0.046 | 55.347 | 50.380 | 2.522 |
| 1200 | 0.047 | 55.348 | 50.381 | 2.523 |
| 2000 | 0.047 | 55.351 | 50.378 | 2.522 |
| grand mean | 0.047 | 55.349 | 50.380 | 2.522 |

Given that the simulated size effect was in line with our empirical results, we next sought to understand the mechanisms by which size reduces error. We evaluated multiple possibilities, none of which were mutually exclusive. One possibility is that the finer tiling of the stimulus space by units in the larger networks allowed the memorandum to be maintained with more precision. This was not the case; networks in which the tuning of the units was matched to the tuning in the 400-unit network and duplicated *k* times to reach a given network size performed comparably to the networks with finer coding (Figure 5A, dotted line).

Another possibility is that error decreases as network size increases due to there being more units from which to decode, allowing the decoder to further average out noise. To address this possibility, we decoded from only 400 units across network sizes, selecting the units matched in tuning to the 400-unit network. While decoding from fewer units increased the decoded error at larger sizes, it did not eliminate the size effect (Figure 5A, dashed line). Furthermore, the benefit of decoding from more units was largely eliminated by averaging the signed error (i.e., decoded position relative to the stimulus) across time (Figure 5 - Supplement 1A); effects of decoding noise may thus not be major drivers of empirical size effects if downstream regions reading out from populations storing the memory integrate the signal over a brief time window, which has been shown to occur (Sharma et al., 2025; Shushruth et al., 2022; van Ede & Nobre, 2024).

### Pure size effects are determined by network-level noise structure

We next asked whether size-dependent changes in the signal and noise characteristics of the networks could have caused the changes in error. We characterized the mean bump profile of the networks (i.e., the signal) by fitting generalized Gaussian functions to network activity across the delay. Confirming the effectiveness of our design, the bump profile averaged across time points was invariant to size (Figure 5 - Supplement 1B, top). After removing the mean activity estimated by the bump fits, we found that the unit-by-unit noise variance was also size invariant (Figure 5 - Supplement 1B, bottom). Together, the size invariance of the unit-level signal and noise characteristics of the network indicate that reduced error with size was a network-level phenomenon.

The attractor encoding the memory occupies a low-dimensional ring in the higher-dimensional space of the network, and noise must be parallel to the attractor if it is to cause the bump position to drift (Figure 5C; (Burak & Fiete, 2012; Wu et al., 2008). Furthermore, random vectors in high-dimensional space tend toward orthogonality as the dimensionality increases (Vershynin, 2018). We therefore hypothesized that as network size —and thus dimensionality—increases, a smaller proportion of the noise relative to the total noise in the network falls along the attractor, leading to less drift in the bump and equivalently less error. To test this possibility, we used the bump fits to characterize the tuning curves for each unit in the network. We then computed a locally-linear approximation to the attractor by taking the derivative of the tuning curves with respect to the initial stimulus, which gives an axis in activity space tangent to the attractor (Figure 5C), analogous to the signal axis in studies of neural activity spaces during perception (Heller & David, 2022; Kobak et al., 2019; Moreno-Bote et al., 2014). We then found the proportion of noise variance out of the total that falls along this axis (see Methods).

As predicted, the proportion of noise variance along the attractor decreased as network size increased. While the proportion of noise variance along the attractor was small at all network sizes, owing to the network connectivity structure it was more than expected for a random vector (Figure 5D; all *p*s = 0.0001), providing evidence that our method identified an important axis in the activity space. Bump attractor dynamics can be approximated by a simple diffusion process in which the variance of the bump position grows linearly with time (Burak & Fiete, 2012; Kilpatrick & Ermentrout, 2013; Wu et al., 2008). Therefore, if the proportion of noise variance falling along the attractor largely or fully explains size-related differences in error, then there should be a strong linear relationship between the proportion of noise variance along the attractor and error variance at the end of the delay. Remarkably, the proportion of variance along the attractor was almost perfectly correlated with the error variance (Figure 5E, *r* = 0.996). That is, given the size invariance of the other properties of the networks, the diffusion coefficient is proportional to the proportion of noise variance along the attractor, establishing a pure size effect on working memory precision.

## DISCUSSION

Despite the critical role of working memory in cognitive functioning and extensively documented individual differences in working memory ability (Barrett et al., 2004; Luck & Vogel, 2013), our understanding of the neurobiological substrates of these individual differences is lacking. Because cortical areas associated with visual working memory are shared with those subserving visual perception (Curtis & D’Esposito, 2003; D’Esposito & Postle, 2015; Serences, 2016), we inferred that they may also share properties determining individual differences; more specifically, we hypothesized that individual variability in the cortical surface area of retinotopically organized visual maps, which is known to impact perceptual performance (Duncan & Boynton, 2003; Himmelberg et al., 2023; Murray et al., 2024; Schwarzkopf et al., 2011; Song et al., 2013), would also impact working memory precision. Our results supported this hypothesis and demonstrated specificity—only the surface area of V1 and IPS2/3 correlated with memory. Furthermore, our neural network modeling identified a novel role of size in determining performance, whereby increases in size reduce the extent to which noise in the network can cause the stored memory to drift. These findings establish a set of neural substrates and mechanisms that together provide a new answer to the critical question of why working memory varies across individuals, which has lingered unanswered despite decades of research devoted to this variability and its impacts..

V1 and IPS have each been frequently associated with working memory storage. Memory contents are decodable from both regions (Christophel et al., 2012, 2018; Ester et al., 2015; Hallenbeck et al., 2021; Harrison & Tong, 2009; Li et al., 2021, 2025; Master et al., 2024; Serences et al., 2009; Sprague et al., 2014; Yu & Shim, 2017), and the quality of these decoded contents correlates with behavior, though with some variability across studies (Emrich et al., 2013; Ester et al., 2013; Galeano Weber et al., 2016; Hallenbeck et al., 2021; Iamshchinina et al., 2021; Li et al., 2021). TMS to either V1 or IPS2 disrupts memory-guided saccade performance (Dake & Curtis, 2025; Mackey & Curtis, 2017), as do lesions to IPS (Mackey, Devinsky, Doyle, Golfinos, et al., 2016), providing evidence for a causal role of both areas. While the relative necessity of each area to working memory storage has been debated (Bettencourt & Xu, 2016a; Hallenbeck et al., 2021; Iamshchinina et al., 2021; Lorenc & Sreenivasan, 2021; Rademaker et al., 2019; Scimeca et al., 2018; Teng & Postle, 2021; Xu, 2017, 2020), our findings add to the preponderance of evidence that both regions are important. One argument for the relevance of V1 to memory storage is its ability to support fine-grained memoranda via its small receptive fields (Christophel et al., 2017), consonant with our finding that spatial working memory precision correlates with the size of V1. Given its substantially larger receptive fields (Mackey et al., 2017), it may seem surprising that IPS2/3 was also correlated with fine-grained spatial precision. However, spatial decoding error in IPS2/3 can be similar to that in visual cortex (Li et al., 2021, 2025). Furthermore, larger receptive fields do not always entail worse decoding (Seung & Sompolinsky, 1993) and can substantially improve the fidelity of neural representations of space (Lehky & Sereno, 2011; Sereno & Lehky, 2011)

IPS2/3 (particularly IPS2) has properties that may make it well-suited to influence spatial working memory storage and behavior apart from mere spatial precision. Persistently elevated neural activity, which is the canonical signature of working memory (Tardiff & Curtis, 2025), in intraparietal sulcus peaks in IPS2/3 (Bray et al., 2015; Hallenbeck et al., 2021; Jerde et al., 2012; Master et al., 2024). These areas are also thought to represent a priority map, which weighs areas of space according to their behavioral relevance (Jerde et al., 2012; Vandenberghe et al., 2012). The size of IPS2/3 could thus correlate with working memory as a source of top-down signals that control the contents of working memory in early visual areas such as V1 (Gazzaley & Nobre, 2012; Jerde et al., 2012; Li et al., 2025; Rahmati et al., 2018; Riddle et al., 2020). IPS2 also contains transformation-invariant object representations, highly overlaps with a functionally defined area of superior IPS that tracks working memory capacity, and has strong saccade-related activity, making it a good candidate for a substrate of working memory storage as well as control (Bettencourt & Xu, 2016b; Silver & Kastner, 2009; Xu, 2018). In contrast, IPS0 may be more involved in lower-level spatial processing than other parts of IPS. Decoding accuracy for spatial working memory peaks in IPS0 and adjacent V3AB (Li et al., 2021, 2025; Master et al., 2024). In keeping with this fine-grained spatial precision, IPS0 demonstrates greater bias for stimuli in the contralateral visual field (Mackey et al., 2017) than other IPS areas and is the only IPS area that overlaps with a functionally-defined region of inferior IPS that is involved in tracking spatial locations (Bettencourt & Xu, 2016b). IPS1 shares many of the characteristics of IPS2, including object invariance, strong responses to saccades, and overlap with working-memory-capacity-related IPS areas (Bettencourt & Xu, 2016b; Silver & Kastner, 2009; Xu, 2018). Based on their functional properties, IPS1 and IPS2 have been proposed as homologues of macaque lateral intraparietal area (LIP; (Silver & Kastner, 2009). However, the retinotopic organization of IPS2 is a better match for LIP (Arcaro et al., 2011), and IPS1 has stronger anatomical connectivity to visual cortex and lower connectivity to frontal cortex compared to IPS2/3 (Bray et al., 2013)

Given our relatively small sample size and the difficulty of interpreting null results, we refrain from drawing strong conclusions about IPS0/1 and other areas that did not correlate with memory. One possibility is that, despite containing decodable working memory information, some of these areas are not causally involved in working memory storage, or at least exert less influence over behavior. It could also be that aspects of their neural architecture make size a less important or unimportant factor. In the case of sPCS, TMS and lesion studies indicate it plays a causal role in spatial working memory (Mackey, Devinsky, Doyle, Meager, et al., 2016; Mackey & Curtis, 2017), possibly by exerting top-down control over visual areas (Comeaux et al., 2023; Curtis & D’Esposito, 2003; D’Esposito & Postle, 2015; Gazzaley & Nobre, 2012; Hallenbeck et al., 2025; Jerde et al., 2012; Li et al., 2025; Merrikhi et al., 2017; Serences, 2016). However, the retinotopic structure of sPCS is less organized than that of visual and parietal cortex, and the adjacent cortex is not retinotopically organized. This means it may be more difficult to accurately estimate the precise boundaries of sPCS using our procedures, which would add noise to any potential relationship between size and memory. It may also be that other neurobiological factors that are known to influence working memory, such as white matter structure (Charlton et al., 2010; Darki & Klingberg, 2015; Vestergaard et al., 2011) or individual differences in the monoaminergic systems (Cools & Arnsten, 2021; Katsuki & Constantinidis, 2012; Mueller et al., 2020; Noudoost & Moore, 2011), are more potent sources of individual differences in this area. Future work with larger samples and additional individual-differences measures should attempt to tease apart these possibilities.

Ultimately, the impact of cortical surface area and other morphological properties on working memory depends on interactions between the type of task and the effect of individual differences in these properties on the underlying neural substrate (Song et al., 2013). Working memory content may be preferentially stored in areas specialized for the perception of those contents (Christophel et al., 2017), and working memory capacity is correlated with gray matter volume of different cortical regions depending on whether the memoranda are spatial locations or abstract shapes (Konstantinou et al., 2017). Even for simple perceptual abilities, interactions between task and neural architecture are crucially important. For example, contrast sensitivity has been shown to correlate with the surface area of V1 when tested with an orientation discrimination task (Himmelberg et al., 2022; Jigo et al., 2023) but not with a contrast discrimination task using white-noise stimuli (Song et al., 2013), potentially because orientation has a topological organization that can be affected by changes in size (i.e., neurons with similar orientation preferences are more proximal and preferentially connected) but contrast does not (Song et al., 2013).

To that end, we designed our study to leverage the retinotopic organization of the dorsal visual stream to tightly connect spatial working memory precision to cortical organization and to our computational model, thereby providing a candidate neural mechanism for size effects. A prior study reported an effect of the size of a region of IPS on spatial working memory capacity (Konstantinou et al., 2017), but measures of capacity do not provide as pure a measure of storage substrates given their additional attentional demands (Fukuda et al., 2015; Shipstead et al., 2014), and the study only used eight equally-spaced spatial positions, which allows for categorical or mnemonic strategies and need not rely on precise spatial memoranda. A study that focused on differentiating neural substrates of capacity from precision also reported that the size of a different region in IPS correlated with working memory precision for orientation (Machizawa et al., 2020), but the study did not use a task that could actually provide a continuous measure of precision, and their design confounded the distinction between capacity and precision with other factors, such as mental effort.

That said, precision and capacity have been argued by others to stem from the same neural resource (Bays et al., 2024), so better understanding the overlap between IPS areas known to correlate with capacity (Todd & Marois, 2004, 2005; Xu & Chun, 2006) and those that correlate with precision could inform these debates. Additionally, neither prior study confined their regions of interest to those with a known cortical organization that could provide a mechanistic basis for size effects. In contrast, Bergmann et al. (2016) found that the size of retinotopically defined V1 predicted orientation working memory, but their task had poor psychometric properties as a measure of capacity and like the Machizawa study did not allow a fine-grained measure of precision, making it difficult to speculate on the basis of the size effect. Therefore, to our knowledge we are the first to report an effect of cortical size on fine-grained memory precision and the first to do so in a way that provides a candidate computational mechanism informed by empirical cortical organization.

There are multiple, not mutually exclusive reasons why the size of a cortical area could affect performance. To model the effect of size, we started from the assumption that increases in size correlate with increases in the number of neurons, which is commonly assumed in size-behavior studies and empirically supported (Kaas, 2000; Marhounová et al., 2019). While prior theoretical (Burak & Fiete, 2012; Zhang, 1996) and computational (Compte et al., 2000) studies have demonstrated the importance of network size on performance in ring attractor models, they did not explain why size should have this effect. We demonstrated that increasing the number of units in the network while holding all other factors constant by imposing scale-invariance was sufficient to improve the memory performance of the network by reducing the drift of the activity bump encoding the memorandum. Importantly, this improvement could be fully explained by proportionally less noise falling along the attractor as the dimensionality of the network increased. This explanation stands in contrast to those offered for size effects in visual perception, which have shown that changes in performance associated with cortical surface area can be explained by changes in neural tuning width (Murray et al., 2024; Song et al., 2013, 2015). That is, given constraints on the physical distance over which local connections between neurons can be made and the topological organization of cortex, greater surface area but fixed connection distances will necessarily result in narrower tuning width in topologically organized cortex. While connectivity constraints exist in cortex (Kaas, 2000; Voges et al., 2010), by relaxing this assumption we showed that changes in tuning width need not be invoked to explain changes in working memory performance with size. Future work should explore how interactions between changes in tuning width and pure size effects affect performance across different working memory domains and tasks.

In summary, our results provide evidence that the size of visual field maps can constrain working memory precision, potentially by influencing the extent to which neural noise can perturb the memorandum. We believe further progress in understanding the neural substrates of working memory and variation therein can be made by carefully combining functional and anatomical neural architecture, precise behavioral assays, and mechanistic models, which in turn could lead to improved methods for supporting and improving working memory function.

## MATERIALS AND METHODS

### Subjects

Twenty-six neurologically healthy human subjects (13 female, 13 male; mean age: 25, range: 20–33) performed the experiment, seventeen of whom were included in a previous report (Hallenbeck et al., 2025). All subjects had normal to corrected-normal vision and were excluded from participation if they had any brain-related medical issues or were currently taking certain drugs (e.g., antidepressants, amphetamines, chemotherapy, etc.). All subjects gave written, informed consent and were compensated $10 per session, with 1–2 sessions per subject.

### Experimental Procedures

Subjects performed a two-item memory-guided saccade task (Figure 2A). Two working memory items (small white dots subtending 0.25°) were presented in the periphery (500 ms). The position of the items was set such that only one item appeared in each hemifield. After a delay period (2,500–3,500 ms, jittered), a response cue displayed at fixation indicated which of the two items was the goal of a memory-guided saccade. Feedback was given by redisplaying the probed item (800 ms) and having subjects make a corrective saccade to this location. Subjects were instructed to maintain fixation during a trial, except when cued to make a saccade. Additionally, a priority cue presented prior to the memory items indicated which item was more likely to be cued for recall. Full task details and the results of the priority manipulation are reported in (Hallenbeck et al., 2025). All analyses in this report collapse across priority condition. Subjects performed 36 trials/run, and completed 11 runs of the task on average (range: 6–20 runs).

### Oculomotor procedures and analysis

Monocular tracking of gaze position was performed with the Eyelink 1000 (SR Research) recorded at 500 Hz. A 9-point calibration routine was performed at the start of each run. If, after multiple attempts, 9-point calibration failed, 5-point calibration was performed.

We preprocessed raw gaze data using custom software (iEye, https://github.com/clayspacelab/iEye). This software implements an automated procedure to remove blinks, smooth the data (Gaussian kernel, 5 ms SD), and drift correct and calibrate each trial using epochs when it is known the eye is at fixation (delay) or the true item location (feedback). Memory-guided saccades were identified during the response period using a velocity threshold of 30°/s. We employed strict data exclusion criteria to ensure errors primarily reflected memory error and not lapses or issues with eye tracking. Trials were flagged for exclusion based on the following criteria: broken fixation during the delay; identified saccade < 2° in amplitude or > 150 ms in duration; RT < 100 ms or > 700 ms; or saccade error > 10°. Runs with fewer than ⅓ of trials remaining after these exclusions were discarded, as this indicated technical problems with the eye tracker or off-task behavior. We further excluded runs in which offline drift correction/calibration indicated outlier variability in head movement (runwise drift correction IQR > 0.927°), calibration failure (< ⅔ of trials with residual calibration error < 1.492°) or failure to make corrective saccades used for offline calibration (> 50% of trials), where the first two thresholds were set as 1.5x IQR above the 75th percentile across subjects. Two subjects had < 125 remaining after these exclusions and were dropped from further analysis. Overall, this resulted in 299 trials on average for each subject (N = 24), with a range of 152–579. We derived memory error from the saccade endpoints by computing the Euclidean distance between the location of the saccade and the true location of the item. We defined the saccade endpoint as the final eye position at the end of the response period, which is thought to lessen the influence of oculomotor factors on the measurement of memory precision (Mackey & Curtis, 2017).

### Magnetic Resonance Imaging

Data were collected at New York University Center for Brain Imaging using a 3T Siemens Prisma MRI scanner (N = 23). Images were acquired using a Siemens 64-channel head/neck radiofrequency coil. Functional volumes were acquired using a T2*-sensitive echo planar imaging pulse sequence with a MB factor of 4 (random dot kinematogram task (RDK): repetition time (TR), 1200 ms; echo time (TE), 36 ms; 56 slices; 2 mm^3^ voxels; rapid serial visual presentation task (RSVP): TR, 1300 ms; TE 42 ms; 56 slices; 2 mm^3^ voxels). High-resolution T1-weighted images (0.8 mm x 0.8 mm x 0.8 mm voxels) were collected at the end of the session, with the same slice prescriptions as for the functional data, and used for registration, segmentation, and display. Multiple distortion scans (TR 6,000 ms; TE 63.4 ms; flip angle, 90°; 56 slices; 2 mm x 2 mm x 2 mm voxels) were collected during each scanning session. The remaining three subjects’ data were acquired using a 3T Siemens Allegra scanner using parameters described in (Mackey et al., 2017).

### Population receptive field mapping and ROI definition

To define the visual field map ROIs, each subject underwent retinotopic mapping in the MRI scanner, following established procedures (Mackey et al., 2017). Subjects maintained fixation at the screen center while covertly monitoring a bar aperture sweeping across the screen in discrete steps, oriented vertically or horizontally, depending on whether the sweep originated from the left or right or top or bottom of the screen, respectively. In the RDK task (N = 17), the bar was divided in thirds, with each segment containing a random dot kinematogram (RDK) used in a match-to-sample task. Subjects reported which of the flanking RDKs moved in the same direction as the central RDK. In the RSVP task (N = 9), the bar contained images of six different objects. On each sweep, subjects were asked to report via button press whether a target image was present. The target image was pseudo-randomly chosen for each run and subjects were familiarized with the image prior to the run. Subjects performed 8–12 runs of pRF mapping. Task difficulty was staircased such that accuracy was maintained around 70–80%. See (Kwak & Curtis, 2022) for further task details.

The resulting BOLD time series were fitted with a population receptive field (pRF) model with compressive spatial summation (Dumoulin & Wandell, 2008; Kay et al., 2013). We then identified visual field map ROIs across the dorsal visual stream. First, we visualized polar angle and eccentricity maps on the cortical surface, thresholded to include only voxels for which the pRF model explained > 10% of the variance (> 8% in frontal cortex). We then identified visual field maps on the basis of polar angle reversals, foveal representation, and anatomical landmarks, using established criteria (Mackey et al., 2017). After maps were drawn, two independent cross-checkers (i.e., not the person who drew the map) verified the maps and noted any disagreements with the map boundaries. Disagreements were resolved via discussion with the first author (N.T.) and map boundaries were edited as needed until agreement was reached.

### Anatomical preprocessing

Cortical surfaces were reconstructed using Freesurfer recon-all (Dale et al., 1999; Fischl et al., 1999). Surface area measurements were then made on the white matter surface (Silva et al., 2018; Winkler et al., 2012) for each ROI and the whole cortex, using mris_anatomical_stats.

### Statistical analysis

Analyses of the behavioral and brain data were performed in Matlab (Mathworks) and R (R Core Team, 2023). Relationships between ROIs or between ROIs and memory error were assessed using Spearman’s rank correlation; semi-partial correlations were used when controlling for covariates. All statistical testing was done via permutation (10,000 samples) using the *permute* package (https://cran.r-project.org/package=permute). Given the nesting of the data within subject and hemisphere, in analyses testing the relationship between memory error and cortical surface area we performed restricted permutations such that hemisphere identity and pairing were maintained (i.e., each pair of hemispheres were permuted as exchangeability blocks (Winkler et al., 2014)). To maintain exchangeability in the presence of covariates in the semi-partial correlation analyses, we used the Freedman-Lane method (Freedman & Lane, 1983), in which samples are generated by permuting the residuals of the regression of the outcome variable (e.g., memory error) on the covariates (e.g., total cortical surface area). We used FDR correction to adjust *p*-values for multiple comparisons (Benjamini & Hochberg, 1995).

### Working Memory Model

We used a neural network model of working memory based on a well-established spiking ring attractor model (Compte et al., 2000). We implemented a mean-field reduction of the model (i.e., a rate code model; (Wong & Wang, 2006) that has been successfully used to simulate memory and decision-making (Engel et al., 2015; Engel & Wang, 2011). The model is highly recurrent, with slow NMDA-like receptors dominating the network dynamics. The dynamics of each unit, which can be thought of as modeling a pool of identically-tuned units in a spiking model, is given by NMDA synaptic gating variable *s,* where:

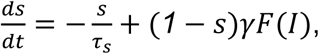

with *γ* = *0*.*641* and *τ*_*s*_ = *60 ms*. The firing rate *r* = *F*(*I*) was a function of total synaptic current *I*:

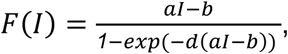

with *a* = *270 Hz*/*nA*, *b* = *108 Hz*, and *d* = *0*.*154 s*. The total synaptic current was the sum of the recurrent current *I*_*r*_, the input stimulus current *I*_*s*_, and noisy background current *I*_*n*_.

The recurrent current to unit *i* was:

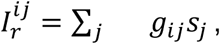

where *g*_*ij*_ was the synaptic weight from unit *j* to unit *i*.

The synaptic weights were structured to provide local excitation based on tuning similarity along with broad, unstructured feedback inhibition. The synaptic weights to each unit *i* with preferred angle *θ*_*i*_ followed a Gaussian profile:

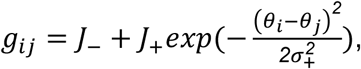

where *θ*_*i*_ − *θ*_*j*_ is the circular difference between the preferred angles of units *i* and *j*, *J*_+_ and *σ*_+_ set the amplitude and standard deviation of the excitatory component of the weights, and *J*_−_ sets the feedback inhibition component (*J*_+_ = *3*.*0 nA*, *σ*_+_ = *32*^∘^, *J*_−_ = −*0*.*546 nA*). To maintain size-invariant bump-attractor dynamics, the weights were then normalized by dividing by the total number of units in the network, *N*. For the primary analyses, unit tuning was distributed evenly in circular space, such that the granularity of the tuning increased with *N*. To account for potential effects of tuning granularity, we also conducted alternative simulations in which networks were constructed by duplicating each unit in the 400-unit network *k* times to achieve a given size (i.e., *k* = *N*/400).

Input stimulus current 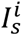 for a stimulus at angle *θ_s_* also followed a Gaussian profile:

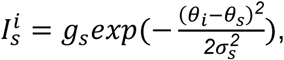

where *g*_*s*_ = *0*.*08 nA* and *σ*_*s*_ = *32*^∘^.

The background current 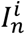 for each unit was a Gaussian white noise process with mean *I*_0_ and standard deviation *σ*_*n*_, filtered by synaptic time constant *τ*_*n*_. Specifically, 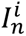 evolved as:

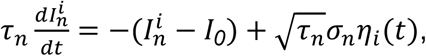

where *τ*_*n*_ = *2 ms*, *I_0_* = *0*.*3297 nA*, *σ*_*n*_ = *0*.*05 nA*, and *η*_*i*_(*t*) was Gaussian white noise with zero mean and unit variance.

Each run of the network simulated a single trial of an angle working memory task. After a 400 ms pre-stimulus period, the stimulus was input to the network for 600 ms. Because the network was translation invariant, we set *θ*_*s*_ = *0*^∘^ on every trial to simplify the analyses. Following stimulus offset, there was a 5 s delay period during which the network maintained a memory of the stimulus in noisy persistent activity. The simulations were implemented in Python. The differential equations were integrated using the Euler-Maruyama method of package sdeint (Aburn, 2017) with a time step of 1 ms.

### Network analyses

To compute the population representation of the memorandum and track error in this representation, we used the population vector (Compte et al., 2000; Georgopoulos et al., 1986; Seung & Sompolinsky, 1993), which is derived from an average of the tuning (in complex polar form) of each unit in the network, weighed by their firing rates:

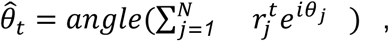

where the decoded angle 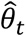 at time *t* is the angle of the vector in the complex plane. We computed the population vector at every time step. To characterize the overall error for each run of the network, we averaged the unsigned error across the last 50 ms of the delay. This measure of error contains variance from two sources: the drift of the bump over time and the noise at each time step. Decoding from more units should reduce the contribution of noise; to quantify this reduction in decoding noise with size, we also decoded the bump position using only the subset of units with tuning corresponding to those in the 400-unit network. To mitigate the influence of decoding noise, in some analyses we instead averaged the signed decoded error over the last 50 ms of the delay and then took the absolute value, which served to mitigate size-related differences in decoding noise.

To better quantify the bump of activity itself, we fit a generalized Gaussian to the firing rates:

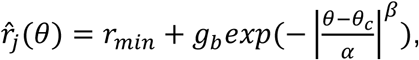

where *r*_*min*_ is the minimum firing rate, *θ*_*c*_ is the center of the bump, *α* controls the width of the bump, *β* modulates the peakedness of the bump, and *g*_*b*_ is a gain term. The bump was fit to activity averaged in 50 ms, non-overlapping time windows, using nonlinear least-squares implemented with scipy.optimize.curve_fit. We restricted our analyses of the bump fits to times > 2 *s* to allow the bump to reach a steady state post stimulus offset. To make fits equivalent across network sizes, for *N* > 400 we only fit the subset of units corresponding to those in the 400-unit network, as in the decoding subset analysis.

#### Noise analysis

Drift in the position of the bump in a ring attractor is caused by noise parallel to the attractor (Burak & Fiete, 2012; Wu et al., 2008). To quantify this noise, we identified the axis in neural space tangent to the attractor—equivalent to the signal axis in studies of encoding and decoding—, which is found by taking the derivative of the tuning curves with respect to the stimulus (Moreno-Bote et al., 2014). Because of the translation invariance of the network, the bump also gives the tuning curve of a unit tuned to *θ*_*c*_ for a stimulus at position *θ*. Therefore, we averaged all fit bump parameters but *θ*_*c*_ across time and network sizes and used those values (Supplementary Table 1) to parameterize the derivative of the generalized Gaussian to derive the signal axis 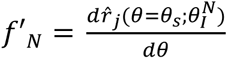, where 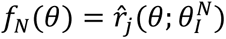 gives the firing rate for each unit for a bump at position *θ*, *θ*_*s*_ = 0 as described above, and 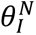 is the vector of unit tunings used to construct the network connectivity for network size *N*.

To quantify the noise structure of each network size, we computed the noise covariance matrix *Σ*_*N*_. To do so, we removed the mean signal at each time point, which was found as the generalized Gaussian 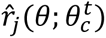, where 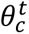 was the fit bump center in the time window containing time *t* and the averaged fit parameters were used for the remaining parameters. We then computed *Σ*_*N*_ from the residualized time series, downsampled by a factor of 5, across all units and simulation runs at each network size. A linear approximation to the proportion of the total noise in the network that falls along the attractor is then given by:

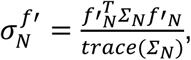

where 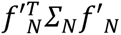 is the noise variance along the signal axis (Bishop, 2006), *trace*(*Σ*_*N*_) is the total noise variance, and *f*′_*N*_ was normalized to unit length. To determine whether the proportion of variance along the signal axis was greater than that expected for a random vector, we sampled *N*-dimensional vectors from a standard normal distribution (9999 samples per *N*) and computed 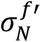 for each sample. *P*-values and 95% confidence intervals were then computed from the null distribution composed of these samples and the empirical 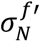.

### Data & Code Accessibility

All analysis and modeling code used in this work is available on a public GitHub repository: https://github.com/clayspace/{TBD}. The functionally-defined regions of interest and behavioral data are available on the Open Science Framework: https://osf.io/{TBD}.

